# TASEP modelling provides a parsimonious explanation for the ability of a single uORF to upregulate downstream ORF translation during the Integrated Stress Response

**DOI:** 10.1101/204784

**Authors:** Dmitry E Andreev, Maxim Arnold, Gary Loughran, Dmitrii Rachinskii, Pavel V Baranov

## Abstract

Translation initiation is the rate limiting step of protein synthesis that is downregulated during Integrated Stress Response (ISR). In our previous work (Andreev, O’Connor et al 2015), we demonstrated that most human mRNAs resistant to this inhibition possess translated uORFs and in some cases a single uORF is sufficient for the resistance. Here we developed a computational model of Initiation Complexes Interference with Elongating Ribosomes (ICIER) to gain insight into the mechanism. We explored the relationship between the flux of scanning ribosomes upstream and downstream of a single uORF depending on uORF features. Paradoxically our analysis predicts that reducing ribosome flux upstream of certain uORFs increases initiation downstream. The model reveals derepression of downstream translation as general mechanism of uORF-mediated stress resistance. It predicts that stress resistance can be achieved with long or slowly translated uORFs that do not favor high levels of translation re-initiation and start with non-leaky initiators.

## INTRODUCTION

In eukaryotes, canonical translation initiation begins with recognition of the m7G cap structure found at the 5’ end of mRNAs. This is achieved by the multi-subunit eIF4F complex consisting of a cap-binding subunit, eIF4E, a helicase eIF4A and a scaffold protein eIF4G. eIF4F then recruits the 40S loaded with eIF2*tRNA*GTP (so called ternary complex, TC), eIF1, eIF1A and eIF5, along with multi-subunit scaffold eIF3 forming a preinitiation complex (PIC). Then the PIC starts to “scan” the mRNA, unwinding mRNA secondary structures and probing mRNA for potential sites of translation initiation. After the initiation codon is recognized, the chain of events leads to large ribosome subunit joining and initiation of polypeptide synthesis. For more details see recent reviews [1–5] on the mechanism of translation initiation in eukaryotes and its regulation.

Not all translation initiation events lead to the synthesis of stable proteins. Many codons recognized as starts of translation occur upstream of annotated coding ORFs (acORFs) encoding functional proteins in many eukaryotic organisms [6–12]. Leader length varies greatly in mammalian mRNAs and at least 20% possess evolutionary conserved AUG triplets upstream of acORFs [13]. The number of potential sites of translation initiation is even higher due to ability of ribosomes to recognize certain near-cognate non-AUG codons (most frequently CUG) as less efficient initiating codons [14–17]. This is particularly prevalent for non-AUG codons located upstream of the first AUG codon, since they are first to be encountered by the PIC [18]. Abundant translation initiation upstream of acORFs has been confirmed by several ribosome profiling experiments [19–21]. Ribosome profiling also revealed that the translation of these ORFs is often altered in response to changes in physiological conditions. A number of stress conditions leads to a global increase in translation of mRNA leaders [22–25]. Sometimes reciprocal changes in acORF translation can be observed among individual mRNAs [22]. One of the stress conditions where uORF-mediated translation control seems to be particularly important is in the ISR [26–28]. ISR involves phosphorylation of the alpha subunit of eIF2 by one of several stress-sensing kinases. This leads to inhibition of its recycling to eIF2*tRNA*GTP carried out by recycling factor eIF2B and to global repression of protein synthesis [29]. We and others recently showed that the translation of a small number of mRNAs are resistant or upregulated during ISR, and the most stress resistant mRNAs possess translated uORFs [26, 30].

The classical mechanism of uORF mediated stress resistance, known as delayed reinitiation, occurs in GCN4 mRNA, the archetypical example of this mechanism in yeast (reviewed in [31]). Although GCN4 regulation involves several uORFs [32–34], only two are absolutely essential for stress resistance. After translation termination at a short uORF located close to the 5’ end, the 40S resumes scanning albeit without the TC. The distance scanned by this ribosome subunit before the ribosome reacquires the TC depends on TC availability. Under normal conditions most of the 40S is quickly reloaded with TC and therefore can reinitiate at a downstream inhibitory uORF. Ribosomes translating this second uORF cannot reinitiate at the acORF start. Under low eIF2 availability, a larger fraction of 40S subunits bypass the second uORF initiation codon before binding of the TC, thereby enabling acORF translation. However, examples, have been found where only a single uORF is sufficient for providing a mRNA with translational stress resistance, e. g. DDIT3 [35].

The start codon of the DDIT3 uORF is in a suboptimal Kozak context and allows for leaky scanning [35, 36]. However, the uORF encodes a specific peptide sequence that stalls ribosomes under normal conditions creating a barrier for trailing PICs which results in strong inhibition of downstream translation [36]. It has been hypothesized that the stringency of start codon recognition is increased during stress and this allows for more leaky scanning [35, 36].

It is also possible that elongating ribosomes translating the uORF obstruct progression of scanning ribosomes downstream and this obstruction is relieved during stress due to reduced initiation at the uORF. Although the obstruction of scanning ribosomes may potentially explain how a single uORF can mediate stress resistance, it is unclear whether such a mechanism is plausible without additional factors involved. While most stress resistant mRNAs possess uORFs, only very few uORF-containing mRNAs are stress resistant [26]. What makes some uORFs permissive for stress resistance? To explore this, we developed a simple stochastic model of Initiation Complexes Interference with Elongating Ribosomes (ICIER) that is based on the Totally Asymmetric Simple Exclusion Process (TASEP). TASEP is a dynamic system of unidirectional particle movement through a one-dimensional lattice, where each site can be occupied by no more than one particle and the probability of particle transition from one site to another is predefined. TASEP is widely used for modelling various dynamic systems, such as road traffic and is also popular in modelling mRNA translation [37–42]. In ICIER we represented scanning and elongating ribosomes as two different particles with different dynamic properties with the possibility of transformation of one into the other at specific sites. The parameters used for the modelling were based on estimates from experimental quantitative measurements of mRNA translation in eukaryotic systems.

The application of the model has demonstrated that indeed a small subset of specific uORFs (constrained by length and leakiness of their initiation sites) are capable of upregulating translation downstream in response to the reduced rate of PIC assembly at the 5’ end of mRNA, which is observed under ISR. Here we describe the computer simulations based on this model and discuss the implications of our results to understanding naturally occurring uORF-mediated stress resistance.

## RESULTS

### The model of Initiation Complexes Interference with Elongating Ribosomes (ICIER)

The ICIER model and its parameters are illustrated in Figure 1. Each site of the TASEP lattice represents a codon. All sites have the same properties except two that represent the start and the stop codons of the uORF. There are two types of particles, scanning ribosomes (**σ**) and elongating ribosomes (***ε***). Each particle occupies 10 codons in accordance with the predominant mRNA length protected by elongating ribosomes [25, 43, 44]. For simplicity, we model scanning ribosome size to be the same even though scanning complexes have been shown to protect mRNA fragments of different lengths [45]. These ribosomes can move forward by a single unoccupied site with probabilities ***m***_***σ***_ and ***m***_***ε***_. In addition, scanning ribosomes can transform into elongating ribosomes at the start site with a probability ***t***_**σ>ε**_, which may vary from 0 (no initiation) to 1 (non-leaky initiation). At the stop site, elongating ribosomes could transform into scanning ribosomes with a probability ***t***_***ε > σ***_ allowing for reinitiation. The remaining elongating ribosomes terminate (disappear) with the probability 1- ***t***_***ε > σ***_ We hypothesized that scanning ribosomes would dissociate from mRNA when upstream elongating ribosomes collide with them. This hypothesis is based on the observation that scanning ribosomes queue behind elongating ribosomes and each other but the opposite has not been observed [45].

Using ICIER under different parameters we explored how the rate of scanning ribosomes arriving at the end of the lattice ***r***_***out***_ depends on the rate with which scanning ribosomes are loaded at the beginning of the latice ***r***_***in***_. ***r***_***in***_ corresponds to the rate of PIC assembly at the 5’ end of mRNA that depends on TC availabilitywhich is reduced upon eIF2 phosphorylation. ***r***_***out***_ corresponds to the rate of scanning ribosomes arrival to the start of the acORF. In essence, upregulation of acORF translation under stress in terms of our model means an increase in ***r***_***out***_ in response to decrease in ***r***_***in***_.

**Figure 1.**
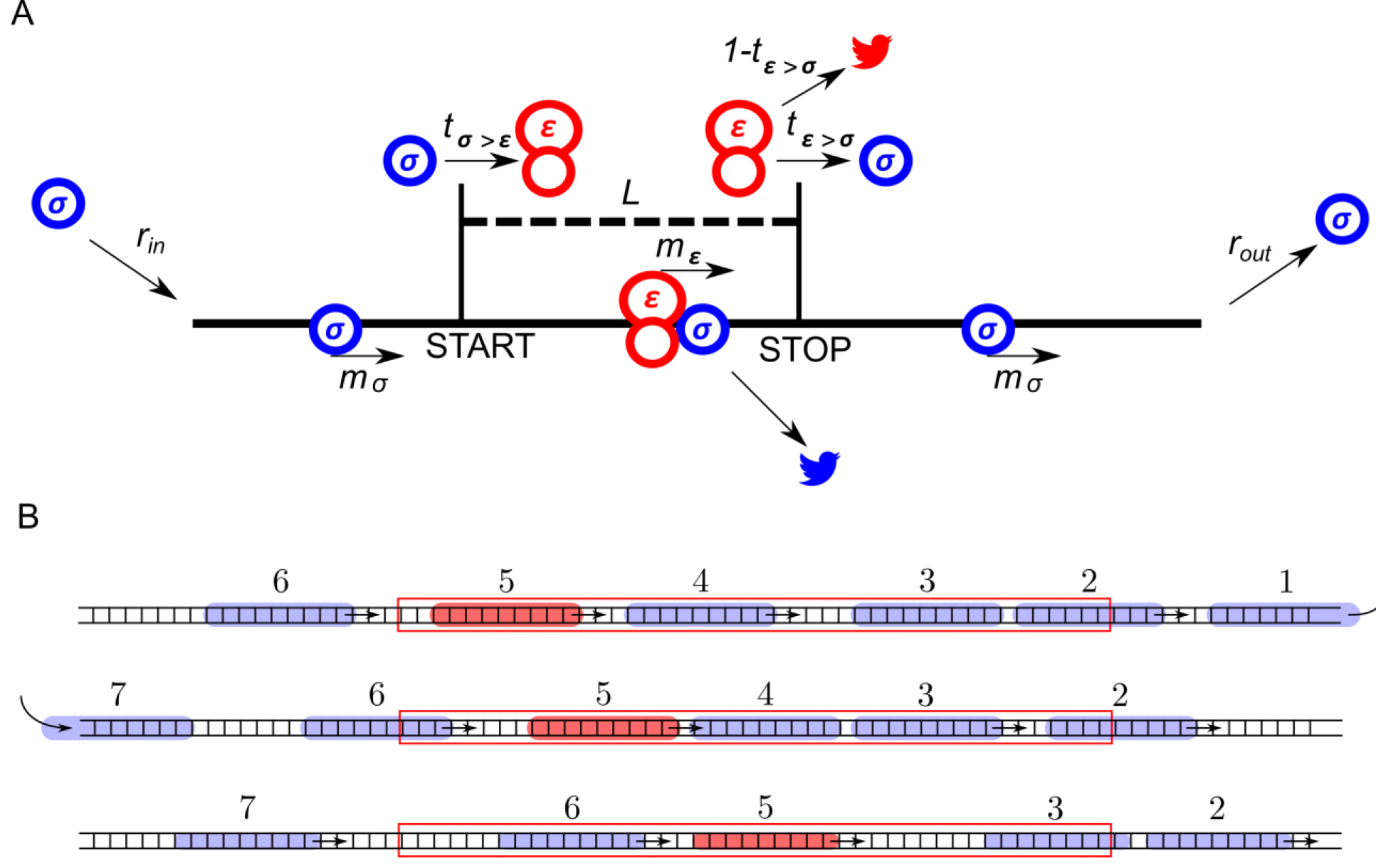
ICIER model. **A.** Parameters of the model. The lattice is shown as a black line with the positions of start and stop codons separated by L sites (codons). Scanning ribosomes are blue and elongating are red in both panels. Ribosome dissociation from mRNAs are indicated with bird symbols. **B.** An example of three subsequent states of the lattice with ribosomes shown as semitransparent oval shapes and each ribosome is labeled with a unique index. Arrows indicate changes between current and subsequent steps. Red rectangle specifies location of the uORF.

### The effect of uORF length on the flux of scanning ribosomes

First, we explored how the rate of PIC loading (***r***_***in***_) effects the density of scanning ribosomes downstream (***r***_***out***_) of a uORF depending on its length (***L***). The results of a typical simulation for a set of specific parameters are shown in Figure 2. For these simulations the possibility of reinitiation was excluded (***t***_**ε > σ**_ = 0). To allow for leaky scanning the strength of uORF translation initiation was set ***t***_**σ>ε**_ = 0.8(80% of scanning ribosomes convert to elongating ribosomes at the start of uORF). The rate of elongation in all simulations was modelled as 0.3 probability that the ribosome moves during a single tact (***m***_**σ**_ = 0.3). Assuming that an average mammalian ribosome moves 5 codons per second during elongation [20], the tact of simulation would correspond to 0.06 seconds (0.3/5). We were unable to find experimental estimates for the velocity of scanning ribosomes*in vivo*, but *in vitro* estimates are similar to that of elongating ribosomes [46, 47]. Hence, for the simulations shown in Figure 2 we used equal rates for elongating and scanning ribosomes (***m***_***ε***_ = 0.3). To simulate stress conditions, we tested the model under variable ***r***_***in***_ from high to absolute zero. It has been estimated that in yeast, on average, the ribosome loads onto mRNA every 0.8 seconds [48], which in terms of our model would be a probability of ribosome load 0.075 per tact. Thus, we decided to model the behavior of the system in the range from 0.1 to 0. In the presence of short uORFs, the ***r***_***out***_ correlates with ***r***_***in***_ although non-linearly (Fig.2A,B black). At the onset of stress, the decrease in ***r***_***in***_ leads to a disproportionally small decrease in ***r***_***out***_ but at a near zero value of rn the rates ***r***_***in***_ and ***r***_***out***_ begin to decrease proportionally. This is because of changes in the likelihood of ribosome collisions that reduce their flow. When the load rate is close to zero, very few ribosomes traverse mRNA making collisions highly unlikely and leading to a direct relationship between ***r***_***in***_. Interestingly, once the uORF reaches a certain length (between 20 and 30 codons in simulations shown in Fig. 2) the relationship between ***r***_***in***_ and ***r***_***out***_ becomes non-monotonous and ***r***_***out***_ starts to increase with decreasing ***r***_***in***_ at a certain interval. The effect is much more profound for longer uORFs (Fig. 2A). However, this advantage afforded by a uORF in stress resistance comes at a price-as can be seen from Figure 2B, uORF containing mRNAs have lower levels of ***r***_***out***_ when ***r***_***in***_ is high and this repression increases with uORF length (Fig. 2D). This occurs due to increasing incidence of collisions by scanning and elongating ribosomes within the uORF․.

**Fig 2.**
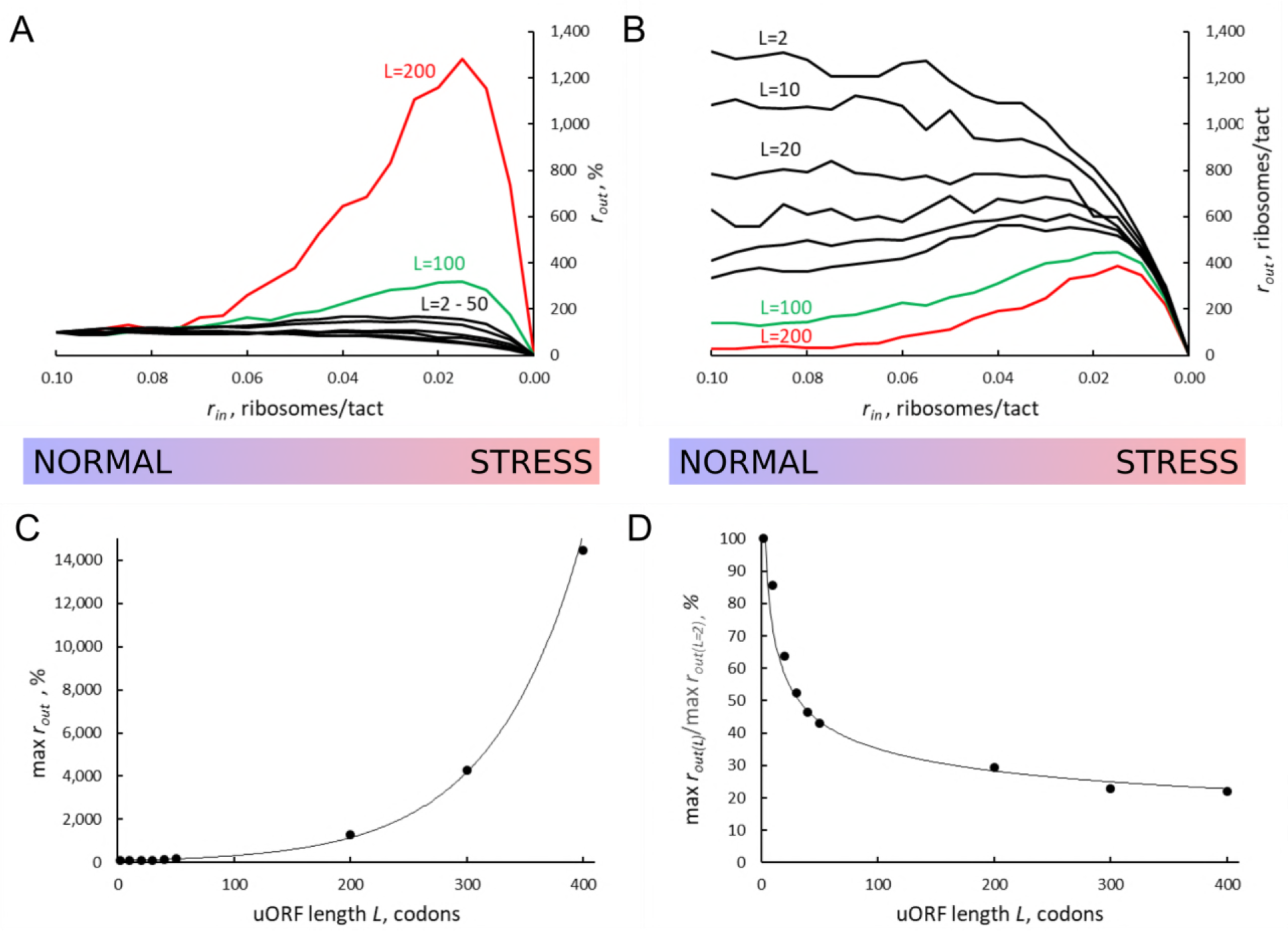
UORF length (***L***) influences how the flux of scanning ribosomes changes downstream (***r***_***out***_) in response to lowered ribosome load (***r***_***in***_) due to stress. Other parameters of the simulations ***t***_**ε > σ**_ = 0 ***t***_**σ>ε**_= 0.8 ***m***_**σ**_ = ***m***_**ε**_= 0.3 **A, B.** Relative (***r***_***out***_(***r***_***in***_)/(***r***_***out***_(***r***_***in***_=0.1)) (A) and absolute (B) changes in the density of ribosomes downstream of a uORF for a range of ***r***_***in***_ from 0.1 to 0 for mRNAs containing uORFs of different lengths (***L***). ***L*** = 2 to 50 are black, ***L*** = 100 are red, ***L*** = 200 are green. **C.** Dependence of the relative (***r***_***out***_) maximum on the uORF length. **D.** The decrease maximum ***r***_***out***_ as a result of increaseing uORF length (maximum ***r***_***out***_ for ***L***=2 codon uORF is taken as 100%).

In other words, according to our ICIER model, long uORFs repress translation of downstream ORFs which are de-repressed during stress. This is consistent with our earlier study, where we found that the best predictor of stress resistant mRNAs is the presence of an efficiently translated uORF combined with very low translation of the downstream acORF [26].

### The effect of uORF elongation rate on the flux of scanning ribosomes

In the simulations described above the movement rates of scanning and elongating ribosomes were set to be equal. However, codon decoding rates can vary significantly depending on their identity and surrounding sequences (for example, see [49–51). Thus, the average elongation rate could also vary for different uORFs. This is particularly salient where uORFs encode stalling sites such as in *DDIT3* [36]. Therefore, we explored how variations in the rate of ribosome elongation affect the behaviour of our model. Even a small decrease in the elongation rate (***m***_**ε**_) strongly increases the maximum ***r***_***out***_ relative to its basal level (Fig. 3). The global (sequence non-specific) rate of elongation may be decreased by stress (for example, by eEF2 phosphorylation) contributing to the stress resistance granted by uORFs. Perhaps, more importantly, the local decoding rates may be specifically tuned to the uORF sequences, through nascent peptide mediated effects or via RNA secondary structures in 5’ leaders.

**Figure 3.**
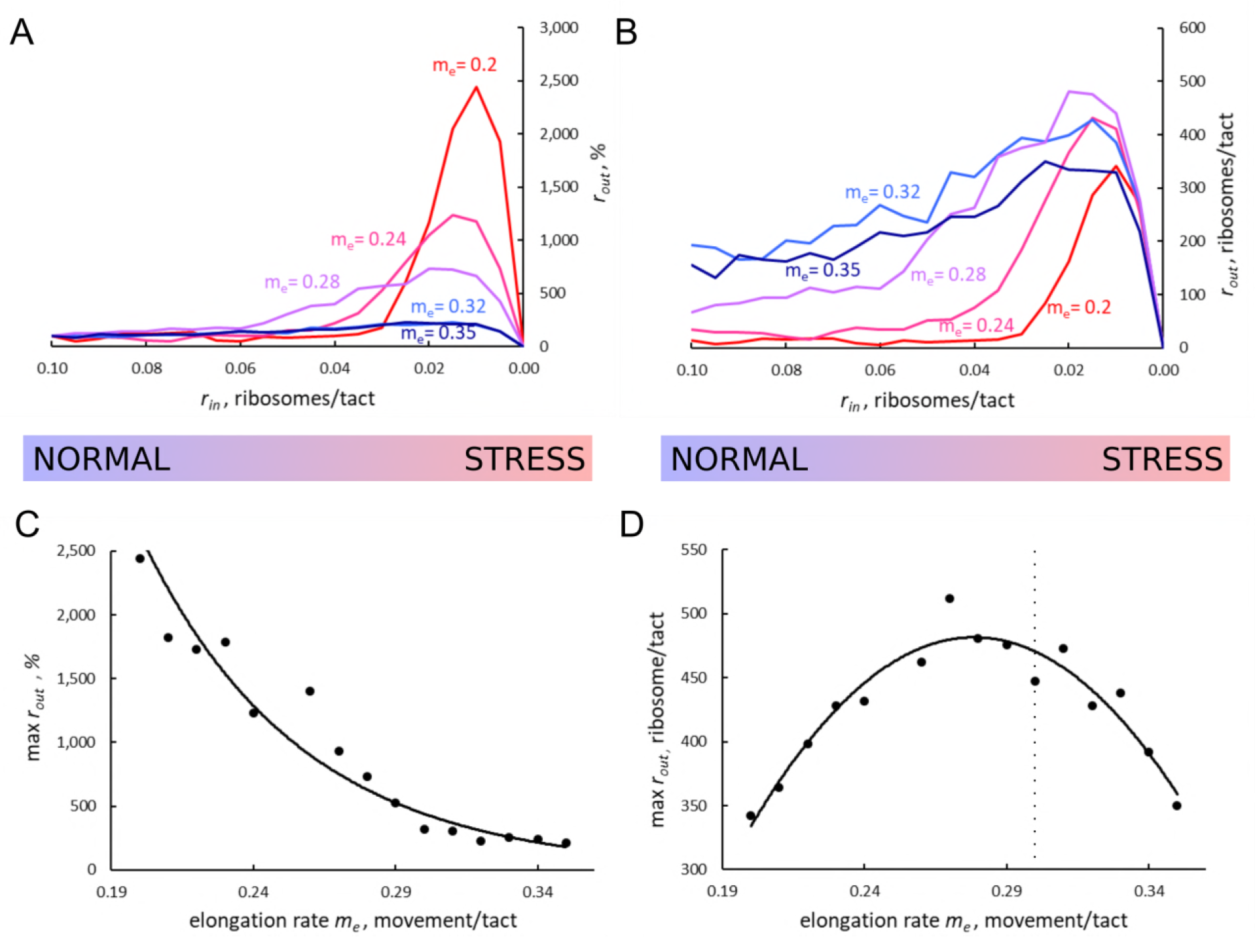
The effect of elongating ribosomes movement rates (***m***_**ε**_) on stress resistance. Other parameters of the simulations ***t***_**ε > σ**_ = 0; ***t***_**σ>ε**_ = 0.8; ***L*** = 100; ***m***_**σ**_ = 0.3. **A, B.** Relative (***r***_***out***_(***r***_***in***_)/(***r***_***out***_(***r***_***in***_ = 0.1)) (A) and absolute (B) changes in the density of ribosomes downstream of the uORF for a range of ***r***_***in***_ from 0.1 to 0 for different elongating ribosome movement rates (***m***_**ε**_ are indicated). **C.** Dependence of the relative (***r***_***out***_) maximum increase on elongating ribosome movement ***m***_**ε**_. **D.** Dependence of the absolute maximum ***r***_***out***_ rate on the ***m***_**ε**_. The dashed line indicates where ***m***_**ε**_ = ***m***_**σ**_.

Whereas the stress resistance monotonously increases with decreased elongation rates (Fig. 3C), the absolute ***r***_***out***_ maximum is not monotonous and reaches a maximum when elongating ribosomes are slightly slower than scanning ones (Fig. 3D). This clearly points to a tradeoff similar to that observed for uORF lengths: The more slowly decoded uORFs provide higher resistance to the stress, but at a cost of higher uORF-mediated repression under normal conditions, thus requiring more mRNA molecules for the same protein synthesis rate. However, if scanning ribosome speed greatly exceeds that of elongating ribosomes, the increased inhibition under normal conditions does not convert into stress resistance. Relative r_out_, as well as absolute r_out_ rates drop with increased elongating ribosome velocities once they approach the velocity of scanning ribosomes (Fig. 3A & B). We do not know whether this observation is relevant to the real world as we do not know if such a situation occurs naturally.

### Strength of translation initiation at uORF and probability of re-initiation downstream

Translation initiation could often be leaky (depending on the context), when only a proportion of ribosomes initiate at a start codon, while the remaining ribosomes continue scanning [2, 52]. This is particularly relevant to uORFs, since, due to their proximity to 5′ ends, even very weak start codons yield detectable translation [18]. Thus, many known regulatory uORFs are initiated at non-AUG starts [15, 22, 27]. As expected, increased leakiness of the uORF start (lower ***t***_**ε > σ**_ elevates the flow of ribosomes downstream of the uORF (Figure 4A). The stress resistance is also reduced considerably for uORFs with leaky starts and disappears when only 30% or less initiate at the uORF in the context of other parameters used for the simulations shown in Figure 4.

**Figure 4.**
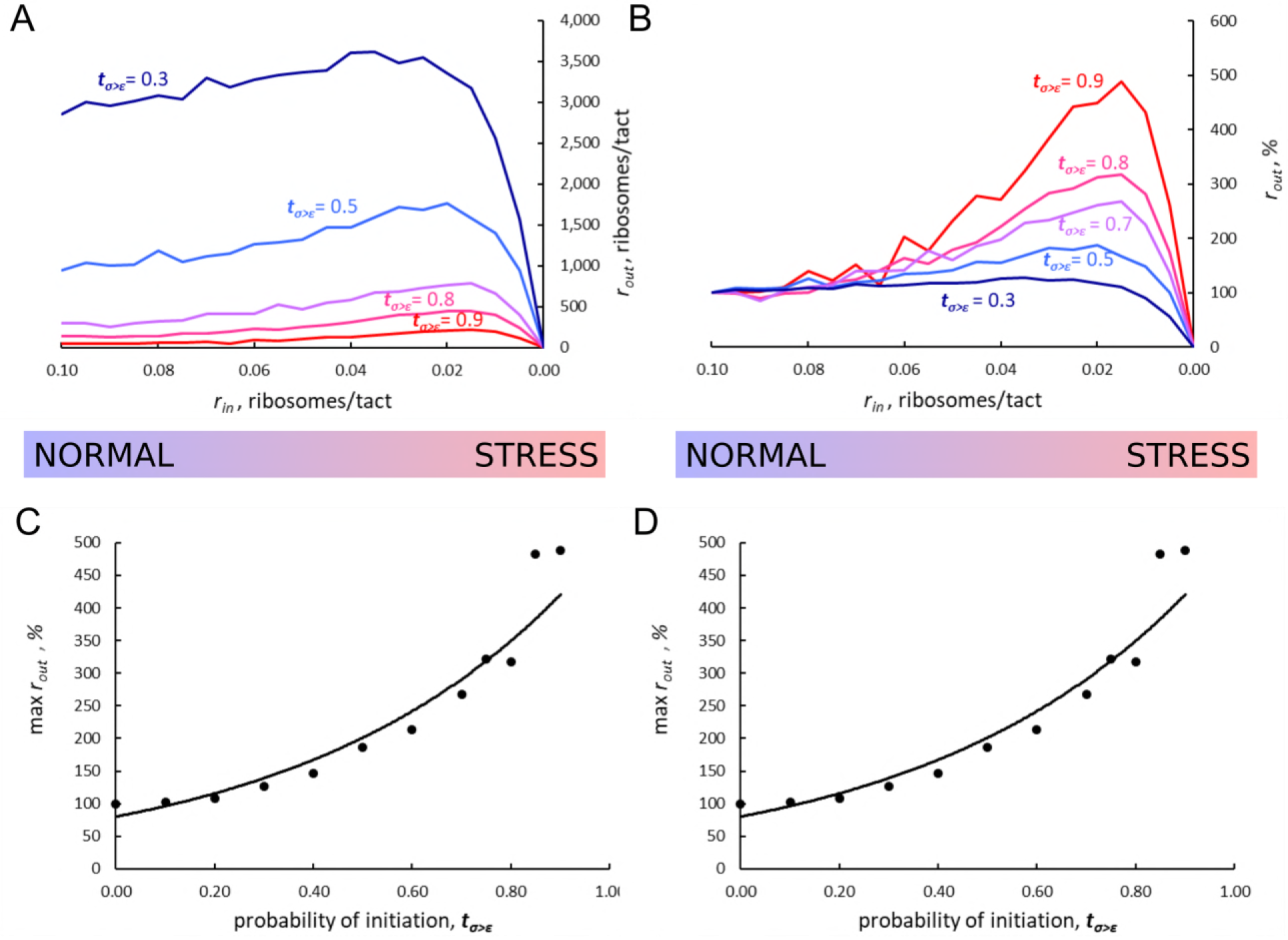
The effect of uORF initiation efficiency (***t***_**σ>ε**_) on stress resistance. Other parameters of the simulations ***t***_**ε>σ**_ = 0; ***L*** = 100; ***m***_**σ**_ = 0.3; ***m***_**ε**_ = 0.3. **A, B.** Absolute (A) and relative (***r***_***out***_(***r***_***in***_)/(***r***_***out***_(***r***_***in***_ = 0.1)) (B) changes in the density of ribosomes downstream of a uORF for a range of ***r***_***in***_ from 0.1 to 0 for different uORF initiation efficiencies (**t_σ>ε_** are indicated). C. Dependence of the relative (***r***_***out***_) maximum increase on the uORF initiation efficiencies ***t***_**σ>ε**_ **D.** Dependence of absolute maximum ***r***_***out***_ rate on the ***t***_**σ>ε**_.

In all of the above simulations reinitiation of ribosomes terminating at the uORF was disallowed, i.e. ***t***_**ε>σ**_ = 0. In practice, however, a significant proportion of ribosomes can reinitiate downstream depending on specific uORF features [32, 53, 54]. We explored how reinitiation alters the behavior of the ICIER model (Figure 5). As expected, elevated reinitiation reduces the inhibitory effect of uORFs and reduces the stress protective effect of uORFs. Interestingly, it appears that a dramatic effect could still be achieved even at very low re-initiation rates. In the simulations shown in Figure 5 the stress resistance almost disappears even when only 5% of ribosomes reinitiate downstream.

**Fig. 5.**
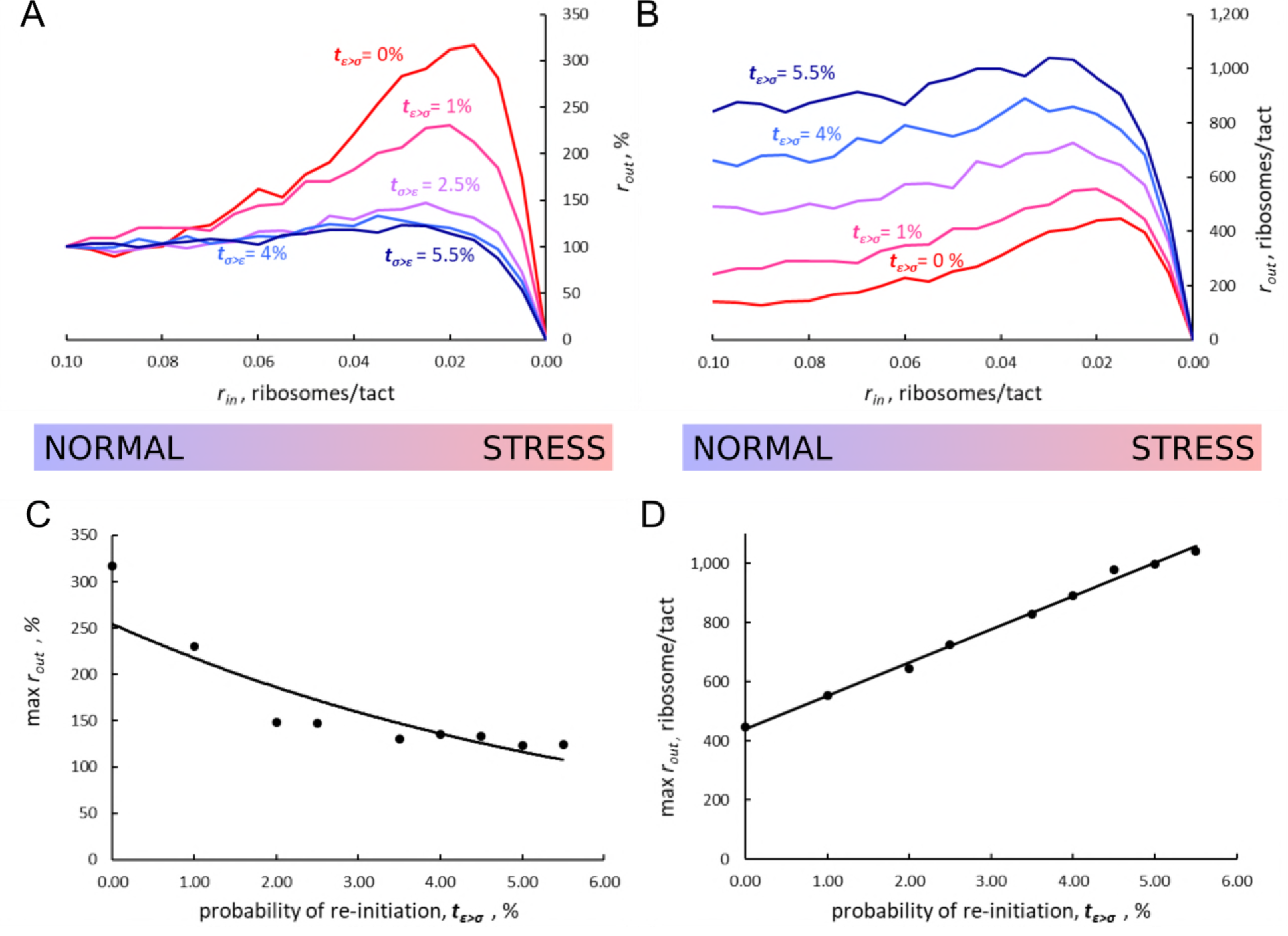
The effect of re-initiation downstream of the uORF on stress resistance. Other parameters of the simulations; ***t***_**σ>ε**_ = 0.8; ***L*** = 100; ***m***_**ε**_ = 0.3; **A, B.** Relative (***r***_***out***_(***r***_***in***_)/***r***_***out***_(***r***_***in***_=0.1)) (A) and absolute (B) changes in the density of ribosomes downstream of a uORF for a range of re-initiation probabilities (***t***_**ε>σ**_ from 0 to 0.055 as indicated). **C.** Dependence of the relative (***r***^***out***^) maximum increase on the re-initiation probability ***t***_**ε>σ**_. **D.** Dependence of absolute maximum ***r***_***out***_ rate on the ***t***_**ε>σ**_.

## DISCUSSION

Using the ICIER model developed in this work we explored how the flow of scanning ribosomes across mRNA is changed in the presence of a translated ORF depending on several parameters such as ORF length, initiation efficiency, decoding rate and re-initiation probability. Our results demonstrate that for mRNAs with certain uORFs the relationship is not monotonous. It appears that within a certain range, the rate of PIC assembly has an inverse relationship with ribosome availability downstream of uORFs. We explored how several different parameters of uORFs affect this phenomenon. It appears that the non-monotonous behavior emerges only after uORFs reach a certain length. The elevation of ribosome flow downstream of a uORF is higher for longer uORFs. However, uORF inhibitory effects on downstream translation also increase with their length. Thus, there is a clear trade-off: while longer uORFs provide stronger resistance to the stress, the translation efficiency under normal conditions is lower for ORFs downstream of longer uORFs. Consequently, to achieve the same protein synthesis rate more mRNA copies need to be synthesized. This trade-off likely shapes the specific lengths of regulatory uORFs and the steady state levels of corresponding mRNAs.

Decreased elongation rates increase the stress resistance. A global decrease in elongation rates is expected under certain stress conditions, for example, due to eEF2K-mediated phosphorylation of eEF2 [55] or due to decreased concentration of available aminoacylated tRNAs. However, the decoding rates may vary among different uORFs since specific sequences affect the speed of the elongation. Indeed, it seems that at least some stress resistant mRNAs contain stalling sites [36]. According to our simulations, leaky initiation at uORF starts reduces stress resistance, as well as the possibility of re-initiation downstream of the uORF.

In all of the above simulations, irrespective of the specific parameters used, the stress resistance (when observed) occurred within a specific narrow range of scanning ribosome loading on mRNA (***r***_***in***_ ~ 0.2). It is reasonable to expect that stress resistance of specific mRNAs would be tuned to particular levels. We equated stress levels to ***r***_***in***_ rates, but it is likely that different mRNAs have different ***r***_***in***_ rates under the same cellular conditions. The exact sequence at the 5’ end of mRNA may potentially influence eIF4F assembly [56] and subsequently the rate with which the PIC is assembled on mRNA. Further, the rate of PIC arrival to the uORF start codon may also depend on mRNA features preceding it. For example, it is well known that a strong stem loop structure located close to the 5’ end inhibits translation, presumably by forcing scanning ribosomes to dissociate from mRNA [57]. Because of this, the exact relationship between the rate of PIC assembly and the level of stress should vary. We therefore predict that due to these differences, optimum translation of specific stress resistant mRNAs should also vary.

In conclusion, by using a simple model of scanning and elongating ribosome interference we demonstrated that a single uORF can be sufficient for providing mRNA translation control with resistance to eIF2 inhibition. The general principle of uORF mediated stress resistance seems to be based on strong repression of downstream translation under normal conditions which is derepressed during the stress. The exact parameters of uORFs are thus a product of the trade-off between the inhibition of absolute levels of translation and its relative increase during the stress. The uORFs providing the resistance are likely to be longer, their initiation sites are expected to have low level of leakiness, they may contain slowly decoded sites and they should not permit high levels of re-initiation downstream. Given that a single uORF could be such a versatile regulatory element, a combination of regulatory uORFs could generate very complex behaviour. Many eukaryotic mRNAs contain multiple uORFs and their concerted impact should be subject of future experimental and theoretical studies. The ICIER modelling may aid the design of mRNA leaders with desired response to stress.

## ACKNOWLEDGMENTS.

This work was supported by Science Foundation Ireland grant (12/IA/1335) to PVB and Russian Science Foundation grant (RSF16-14-10065) to DEA. DR acknowledges the support of NSF through grant DMS-1413223.

